# *De novo* genome sequence assembly of the RNAi-tractable endosymbiosis model system *Paramecium bursaria* 186b reveals factors shaping intron repertoire

**DOI:** 10.1101/2024.08.09.607295

**Authors:** Guy Leonard, Benjamin H. Jenkins, Fiona R. Savory, Estelle S. Kilias, Finlay Maguire, Varun Varma, David S. Milner, Thomas A. Richards

## Abstract

How two species engage in stable endosymbiosis is a biological quandary. The study of facultative endosymbiotic interactions has emerged as a useful approach to understand how endosymbiotic functions can arise. The ciliate protist *Paramecium bursaria* hosts green algae of the order Chlorellales in a facultative photo-endosymbiosis. We have recently reported RNAi as a tool for understanding gene function in *Paramecium bursaria* 186b, CCAP strain 1660/18 [1]. To complement this work, here we report a highly complete host genome and *trans*criptome sequence dataset, using both Illumina and PacBio sequencing methods to aid genome analysis and to enable the design of RNAi experiments. Our analyses demonstrate *Paramecium bursaria*, like other ciliates such as diverse species of *Paramecia*, possess numerous tiny introns. These data, combined with the alternative genetic code common to ciliates, makes gene identification and annotation challenging. To explore intron evolutionary dynamics further we show that alternative splicing leading to intron retention occurs at a higher frequency among the smaller number of longer introns, identifying a source of selection against longer introns. These data will aid the investigation of genome evolution in the *Paramecia* and provide additional source data for the exploration of endosymbiotic functions.

## Introduction

Endosymbiosis is a key phenomenon which has played an important role in the early evolution of eukaryotic cellular complexity [2] and the diversification of eukaryotic forms from algae to corals to insects (e.g., [3–7]). *Paramecium bursaria* (*Pb*) is a ciliate protist and a member of the Alveolata supergroup [8]. Like all ciliates, *Pb* possesses two nuclei: a macronucleus which encodes somatic function and is typically characterised by short chromosomes with a high ploidy count, and a *trans*criptionally inactive diploid micronucleus which engages in infrequent sexual reproduction [9]. This single-celled organism in its natural state hosts in excess of 100 green algae in a stable but facultative endosymbiosis [10]. Cell sampling, culturing and rDNA marker sequencing combined with phylogenetics has shown that the *Pb* species complex is composed of numerous ‘syngens’ (i.e., complementary mating type groups) with variant biogeographical provenance and which may represent cryptic species [11,12].

The *Pb* system has emerged as a powerful model system for conducting experimental research on how two distinct organisms function within an endosymbiotic interaction (e.g., [10,13–16]). As a photosymbiotic protist, *Pb* cultures are easy to grow, and several characteristics of the endosymbiotic interaction are directly observable using microscopy. These characteristics include, for example, the chlorophyll status of *Pb* cells (a proxy for the wider status of the algal population) [17]. Important work has also shown that the endosymbiosis is based on algal secretion of photosynthesis-derived fixed carbon in the form of maltose. This behavior is triggered by moderately acidic pH conditions; a likely outcome induced when the algae become enclosed within the host phagotrophic derived symbiosome, known as the perialgal vacuole [16,18,19]. Much of the experimental progress is underpinned by the capacity to separate host and endosymbiont, culture the partners separately, and then re-initiate the endosymbiosis [16,18]. The capacity to separate the partners has been used to explore compatibility between different strains of host and endosymbiont, demonstrating variant interaction responses [20–24]. This indicates that variant syngens have distinct genetic, phenotypic and endosymbiotic interaction characteristics.

Our long-term aim is to develop *Pb* 186b as a model organism for studying the cell biology of facultative phototrophic endosymbiotic interactions in order to understand how cellular mechanisms that support endosymbiotic interactions evolve and allow for interaction stability. To this end, we and others have developed RNA interference (RNAi) gene knock-down methods which can allow rapid assessment of gene functions which control endosymbiotic interactions [1,17,25]. We found *Pb* 186b (CCAP 1660/18) [11,26,27] is readily tractable for RNAi, and suspect that amended culture conditions would make other strains also tractable. We note that sequencing initiatives have produced draft genome assemblies for five additional strains of *Pb* [14,25,28]. To further facilitate the use of *Pb* 186b for functional experimentation, we report here the draft macronuclear genome assembly and annotation of this RNAi-inducible strain. Genome annotation of ciliates such as *Paramecium* with a macronuclear chromosome structure, non-universal genetic code [29] and tiny introns [30,31] is a significant challenge. To this end, we also report a comparative analysis of intron sequence variation across a subset of *Paramecia* species, demonstrating an intron profile dominated by tiny introns. We use PacBio Iso-Seq analysis to identify alternative splicing in *Pb* 186b, demonstrating that larger introns show a much higher rate of intron retention than the more numerous tiny introns.

## Results & Discussion

### Genome Assembly and Annotation

Here, we have generated genome and *trans*criptome sequencing data using PacBio and Illumina methodologies for *Paramecium bursaria* 186b (CCAP 1660/18). The *Pb* culture represents a consortium of the host ciliate, one or more species of endosymbiotic green algae from the order Chlorellales, candidate bacterial endosymbionts, and bacterial food. We separated the initial read libraries into putative taxonomically distinct bins, three of which contained *Pb* signal. These three bins were combined to form an initial assembly of 1,179 contigs totaling 50.93 Mbp, with an N50 of 96.2 kbp, (where 69.96% of the genome is contained in contigs > 50 kbp). As expected for *Paramecia* species [9,32], this assembly had a low GC content of 26.4% (Figure 1A).

**Figure 1.**
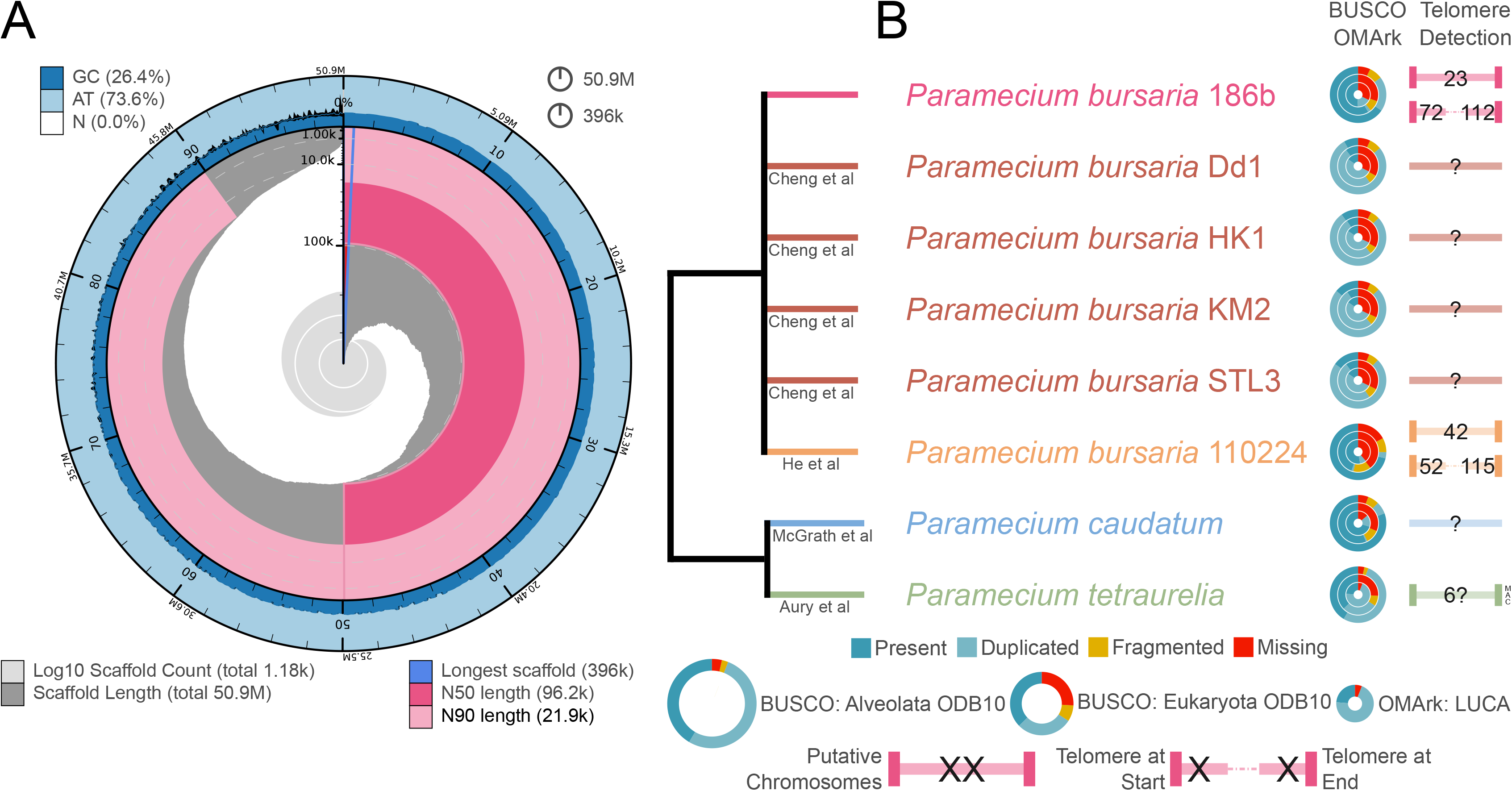
Summary genome assembly statistics and comparison with other *Paramecia* genomes. **A**. A snail diagram from Blobtools demonstrates the summary assembly data for *Paramecium bursaria* 186b, including GC content, overall size (Mbp), number of and the longest scaffold(s) and the N50. Circumferential (50.9 M) and radial scales (396 k) are shown, top right. B. Summary cladogram showing a basic tree topology for three *Paramecia* species. Genome source data includes [9,25,28,32]. Species nodes are labelled with BUSCO and OMArk assembly completion statistics (in concentric rings) and evidence of telomere structures when recovered, see key for further details. Question marks denote no known telomeric sequences have been found or they are unconfirmed in the original publication.

Blob-plot analysis of the initial assembly showed putative contamination with 21 contigs identified as belonging to the Bacteriodetes (*Pedobacter*) or Proteobacteria (*Rickettsia*) groups (0.56 Mbp, ∼1% of the total sequence assembly). Bacteria from these groups have previously been shown to form endosymbiotic associations with other *Paramecia* [33–37], suggesting this contamination was derived from symbiotically associated bacteria, with putative *Pedobacter* and *Rickettsia* contigs seen in the additional read library bins. These 21 contigs were removed from the assembly prior to downstream analyses and gene prediction. A further three contigs with putative mitochondrial signal, and one contig made up entirely of repeat sequences (not a putative telomeric sequence, as identified by manual inspection) were also removed. Further ‘contamination’ in the form of a PacBio blunt-end adapter was present in eight contigs. Four of these contigs were split into two (by removing the adapter sequence), and for the other four the adapter was trimmed from the 3’ ends. This process left a total of 1,158 contigs totaling 50.27 Mbp (L/N50 = 153/96.84 kbp, Table 1).

**Table 1.**
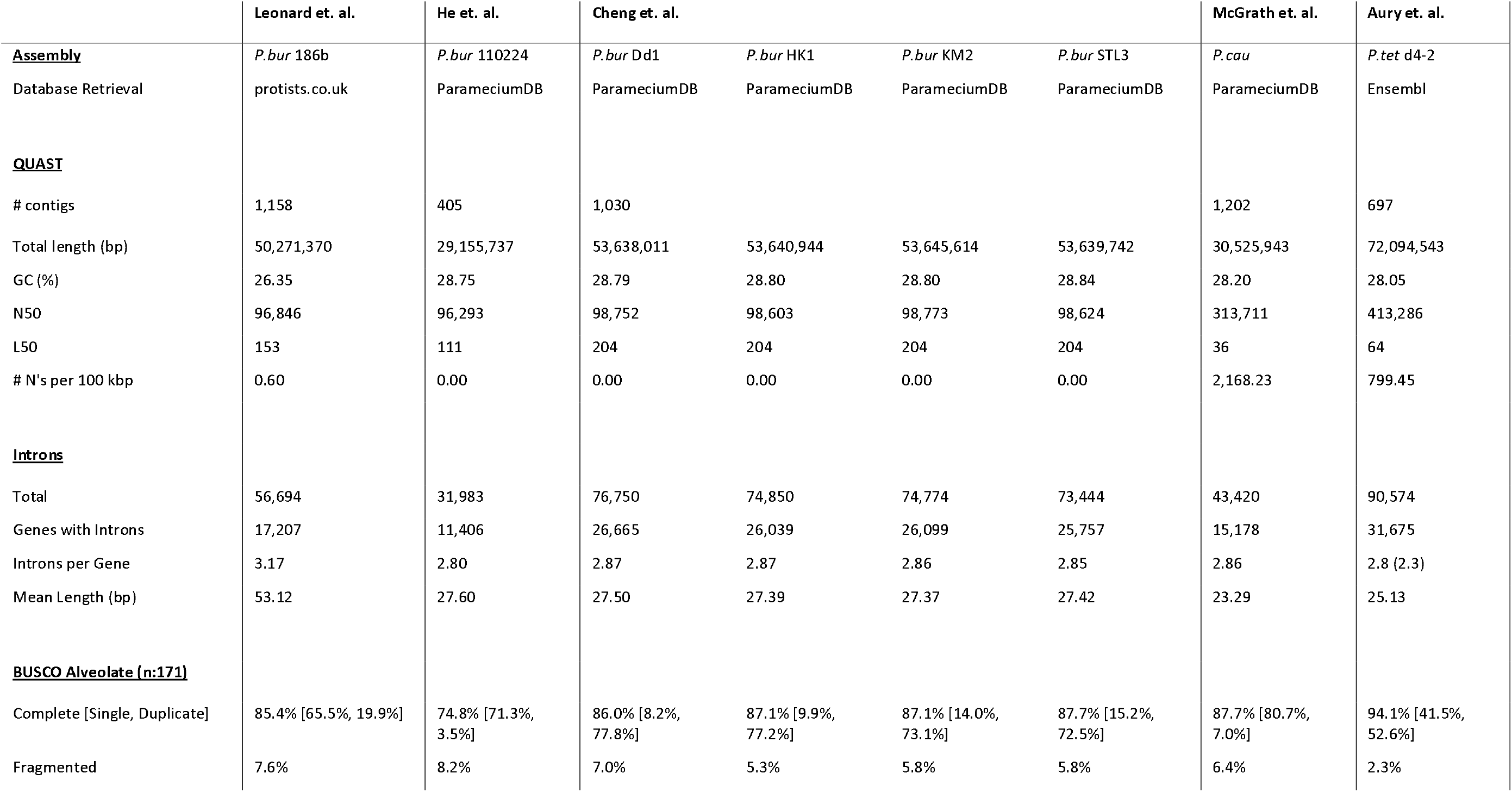
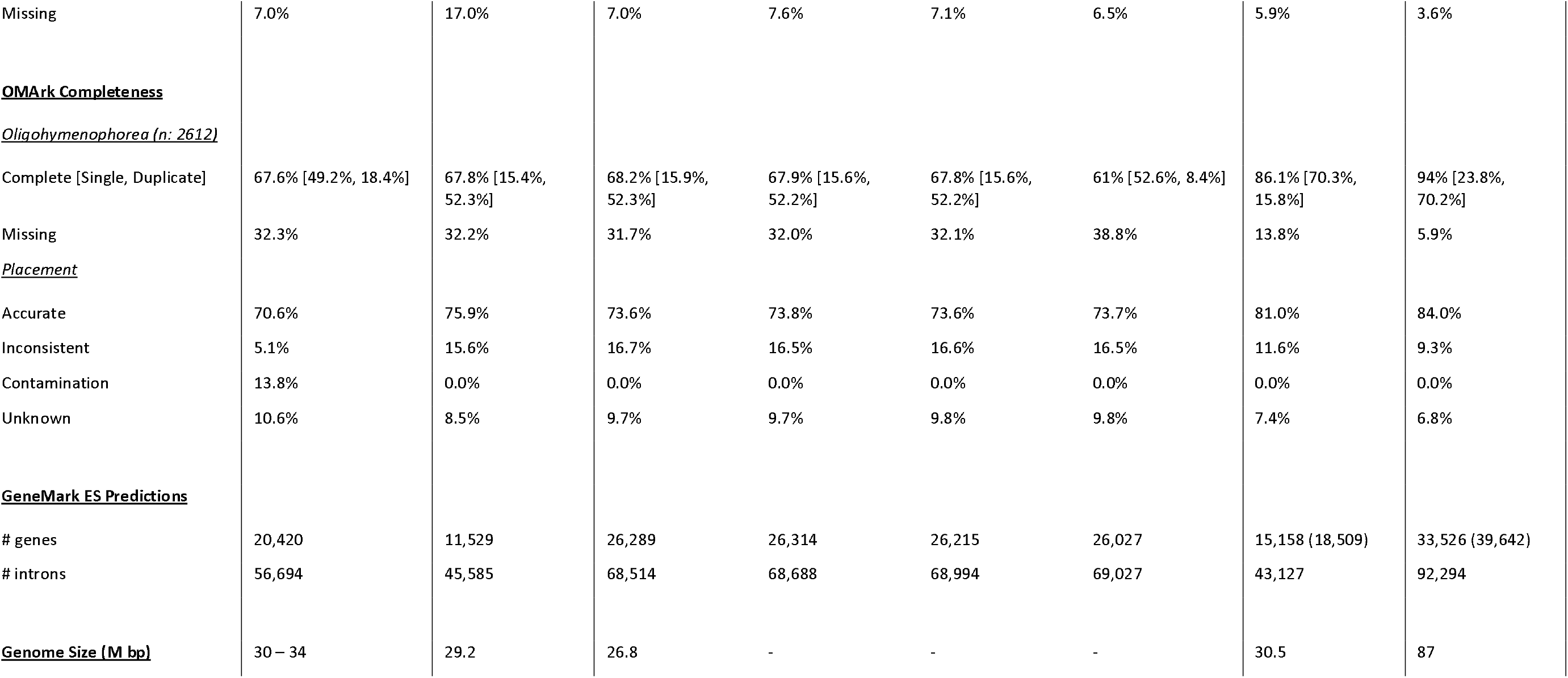
Comparison of genome assembly data. Numbers in round brackets are derived from genome portals, other numbers are from reanalysis (generated by QUAST [89,90]) so comparisons are standardised.

The assembly profile is consistent with the short numerous chromosome structures typical of *Paramecia* macronuclei [9, 28] and shows similar assembly statistics to previously sequenced *P. bursaria* strains across most metrics (Figure 1B and Table 1). Mean coverage of the genome assembly with PacBio CLR was 1,907X σ 1,533X, and Illumina NovaSeq 3,595X σ 3,042X with high levels of variation in coverage here likely a product of the chromosome structures of *Pb* [28]. Genome size estimation using GenomeScope 2.0 [38] and the Illumina NovaSeq data suggests a length between 30 and 34 Mbp, which is comparable to previously reported *P. bursaria* strains (Dd1 = 26.8 Mbp & 110224 = 29.2 Mbp) and *P. caudatum* (Pc) with 30.5 Mbp. Although our reported assembly size is larger than other *Pb* strain genome assemblies, both estimators of genome completeness demonstrate similar levels of completion across multiple *Pb* strain assemblies (Table 1). Specifically, BUSCO [39,40] analyses suggest a relatively complete genome with a score of 60.8% using the Eukaryota ODB10 database and 85.4% using the Alveolate ODB10 database. The program OMArk [41,42] (alignment free proteome completeness assessment using the OMA orthology database) gave a completeness score of 67.6% using the LUCA dataset (with Oligohymenophorea identified as the best taxonomic affiliation). Variances between assembly sizes could be the product of differential inclusion of micronuclear chromosomal sequence within the macronuclear assembly bins across the different genome projects. We note that our assembly contains 115 contigs (0.17 Mbp) with no identifiable genes which may represent fragments of micronuclear chromosomes which are known to be characterised by jumbled ORFs [43]. Alternatively, the differences in genome size estimation could stem from different genome assembly and gene prediction strategies, and/or genuine genome variation between *Pb* strains (as previously suggested [28]).

Ciliate macronuclear genomes are often composed of numerous short chromosomes [28,44,45]. To further investigate chromosome structure in the *Pb* assemblies we searched for identifiable telomere-like sequence tracks using both BLASTn-short and Tapestry searches [46,47] using the previously identified ciliate telomere sequence [45] of “5’-CCCCAACCCCAA-3’” and its reverse complement “TTGGGGTTGGGG”. To test for other repeat motifs which could represent variant telomere structures, we used TelFinder [48] which searches for repeat motifs via k-mer analysis at the ends of scaffolds. Whilst several other repeat motifs were found, none were represented more than once across the scaffolds (at either end) and so they are unlikely to represent any varied, missed or alternative telomere-like structures.

In total we identified 207 (17.9%) contigs with telomere-like sequence motifs; of which 23 (∼2%) contained telomere-like sequence motifs at both ends of the contig, 72 at the 5’ and 112 at the 3’ end. The same analyses were conducted for all five additional publicly available *Pb* genome assemblies [25,28], along with *Paramecium caudatum* (*Pc)* [32] and *Paramecium tetaurelia* (*Pt*) [9] genome sequence datasets, identifying additional ‘complete’ chromosome-like structures in only the *Pb* 110224 and *Pt* assemblies. Considering the contigs with putative telomeres on both ends, these data demonstrate mean chromosome lengths of 34,952 bp (n = 23, median = 25,547 bp) for *Pb* 186b, 88,121 bp (n = 42, median = 79,007 bp) for *Pb* 110224, and 221,083 bp (n = 6, median = 284,010) for *P. tetraurelia* (Figure 1B). These data show high variability in chromosome size but are consistent with *Pb* 186b possessing small macro-nuclear chromosomal structures [28].

To further explore genome completion and to facilitate gene prediction and annotation we generated a PacBio Iso-Seq *trans*criptome library. Gene predictions were completed using a combination of GeneMark-ES v7.4 [49] using the ‘--gcode 6’ option (to account for ciliate stop codon usage), and a modified version of funannotate [50] incorporating the Iso-Seq data and Illumina RNA-Seq library. This produced 20,420 putative genes, with a median size of 1,302 bp (mean of 1,785 bp), a median intron size of 25 bp (mean of 26.2 bp) and a median exon size of 224 bp (mean of 433 bp). The PacBio Iso-Seq refinement and clustering workflow [51] includes a step for mapping the final reads to the genomic scaffolds. This identified 44,631 *P. bursaria* candidate *trans*cripts with 99.9% mapping directly to the genome assembly, with only 37 (0.08%) candidate *trans*cripts that did not map. This result is also consistent with a high level of completion for the macronuclear genome assembly.

### Paramecium intron diversification dynamics

*Paramecia* have been shown to possess a high number of short introns [30]. This feature, combined with the reduced repertoire of stop codons within the ciliate non-standard genetic code [29], means that identifying accurate open reading frames and, therefore, gene models is a significant challenge. Candidate introns were recovered by extracting putative intron sequences from the gene predictions (see Methods). These data demonstrate 56,694 introns with 17,207 genes possessing one or more intron and 13,004 genes with two or more introns.

This is an average of 3.17 introns per gene, a similar statistic to other *Paramecia* when processed using the same bioinformatic pipeline (Table 1). These results also suggest a higher number of introns per gene for *Paramecium tetraurelia* (2.8 introns per gene using the same pipeline) than previously reported ([30,52] – 2.3 introns per gene), but among a smaller number of gene models identified using our gene annotation pipeline (Table 1). Previous work suggests that *Paramecium tetraurelia* possesses a large number of ‘cryptic introns’ [30,52], which may lead to variant estimations of intron number using different bioinformatic methods. We also note that the increased number of introns recovered for *Pb* 186b may stem from the use of Iso-Seq long read *trans*criptome sequencing which is likely to be more sensitive at recovering longer intron forms compared to the RNA-Seq methods used previously.

These data also showed a dominant proportion of 23 bp introns (highlighted in blue in Figure 2) in *Pb* 186b and all other *Pb* genomes. In contrast, *P. caudatum* has a dominant intron size of 22 bp, while *P. tetraurelia* has a dominant intron size of 25 bp [9] (Figure 2). The dominance of 23 bp introns in the *Pb* genomes is notable as it is the same size as the small interfering RNAs (siRNAs) used by the RNAi pathway in both *Pb* and *Pt* [1] to facilitate gene knock-down [53], and endogenous small RNAs (sRNAs) have been shown to regulate expression of both *cis an*d *trans ge*ne targets in *Pt [5*4]. Given the overlap in size of the dominant intron population and sRNA population in *Pb* 186b [1], we tested for the possibility that the *Pb* 23 bp introns are retained and present in the *Pb* sRNA population. To do this we searched the assembled sRNA sequencing data from *Pb* 186b [17] with BLAT (enabling -minScore=15), identifying no evidence that the *Pb* 23 bp introns are detectable among the *Pb* 186b sRNA *trans*cripts. This is consistent with similar analysis from *P. tetraurelia* [47,55], which demonstrated a low rate of recovery for siRNAs containing intron sequences.

**Figure 2.**
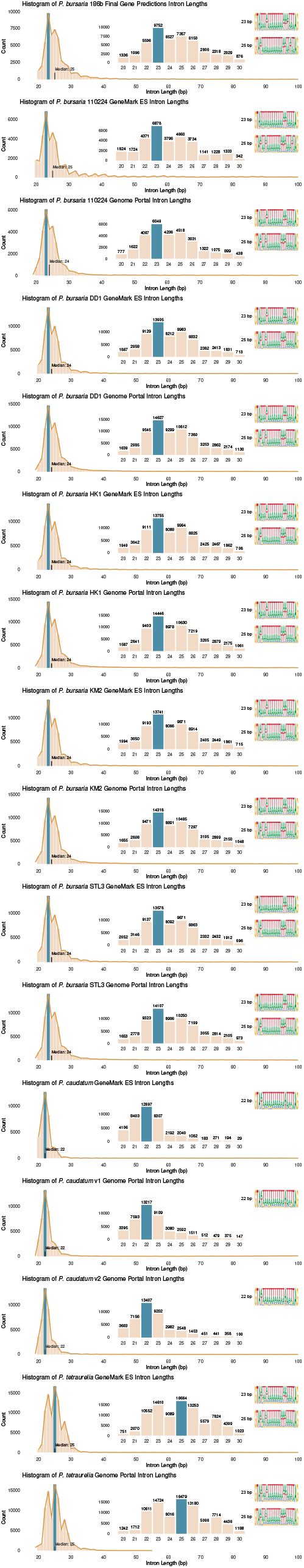
Analysis of intron size distribution and dominant intron sequences for the *Paramecia* species compared. Sequence logos were calculated using [87,88]. Gene predictions for the five other *Pb*, along with *Pc an*d *Pt sp*ecies were recalculated with GeneMark ES [72,73] to provide direct comparisons and appear first in pairs with their genome portal gene predictions. We note that intron predictions have been generated by the use of different sources of *trans*criptome data (*Pb:* Iso-Seq, *Pc & Pt:* RNAseq) and this may have led to an increased recovery of longer-form introns for *Pb an*d therefore increased the median measure of intron size for this species.

To explore intron sequence variation, we calculated sequence logos revealing that the 23 bp introns of *Pb* are largely composed of a core AT rich region and bear a GT-AG splice sites like other *Paramecia* introns (Figure 2) [31]. These sequence logos also show that intron sequences are conserved across the *Pb* species complex (6 genomes), but demonstrate that *Pb* 186b possesses variant AT motifs compared to the other *Pb* strain sequenced (Figure 2), thereby demonstrating an aspect of genome variation among different strains.

### Paramecium bursaria alternative splicing dynamics

The results reported here provide evidence for 20,420 predicted genes with 44,631 distinct Iso-Seq *trans*cripts. This is suggestive of a substantial level of alternative splicing (AS) and is consistent with previous suggestions for other *Paramecia* species [52]. To further explore these data, we detected AS events by mapping the Iso-Seq full length non-concatemer *trans*cripts using the program HISAT2 [56] onto the gene models and then using IsoQuant [57] for isoform discovery. This approach detected 23,871 candidate alternatively spliced events mapping to 4,362 *Pb* genes, demonstrating that ∼22% of the *Pb* 186b gene repertoire shows putative evidence of alternative splicing.

Next, we used the IsoQuant pipeline to identify AS corresponding to putative intron retention (IR) events. IsoQuant identifies three categories of IR: ‘incomplete_intron_retention_3’ which are intron retention events at the 3’ end of the mRNA that led to partial or truncated splice variants (261 events identified); ‘incomplete_intron_retention_5’ which are intron retention events at the 5’ end that led to partial or truncated splice variants (184 events); and standard ‘intron_retention’ splice variants (698 events) where the intron is retained within the core of the mRNA sequence. The IsoQuant pipeline identifies an additional category called ‘fake_micro_intron_retention’, which include annotated introns < 50 bp in length which are described in the IsoQuant manual as artifacts i.e., ‘short annotated introns that are often missed by the aligners’. As *Pb* has short introns, we manually checked alignments and found that they represent true IR events (1,340). In total we identified 2,483 IR events, ∼10% of the total number of 23,871 AS events. These IR events correspond to 2,473 genes (or ∼12% of the gene repertoire).

To explore AS in *Pb* further, we quantified the ratio of IR events per intron length and compared this to the distribution of introns from 20 bp to 100 bp (Figure 3). These data indicate that relative IR rate increases as the intron size increases. We note that the rate is partly an effect of the small number of longer introns present. A Gompertz function was fitted to the data using non-linear least squares to estimate the intron length at which intron retention ratio does not continue to increase (i.e., reaches an asymptote). We computed 95% confidence intervals around the estimated asymptote, and, using the lower confidence bound as a conservative threshold, identified 52 bp as the intron length at which the intron retention ratio reaches the asymptote (represented as a black dotted line on Figure 3 & S1).

**Figure 3.**
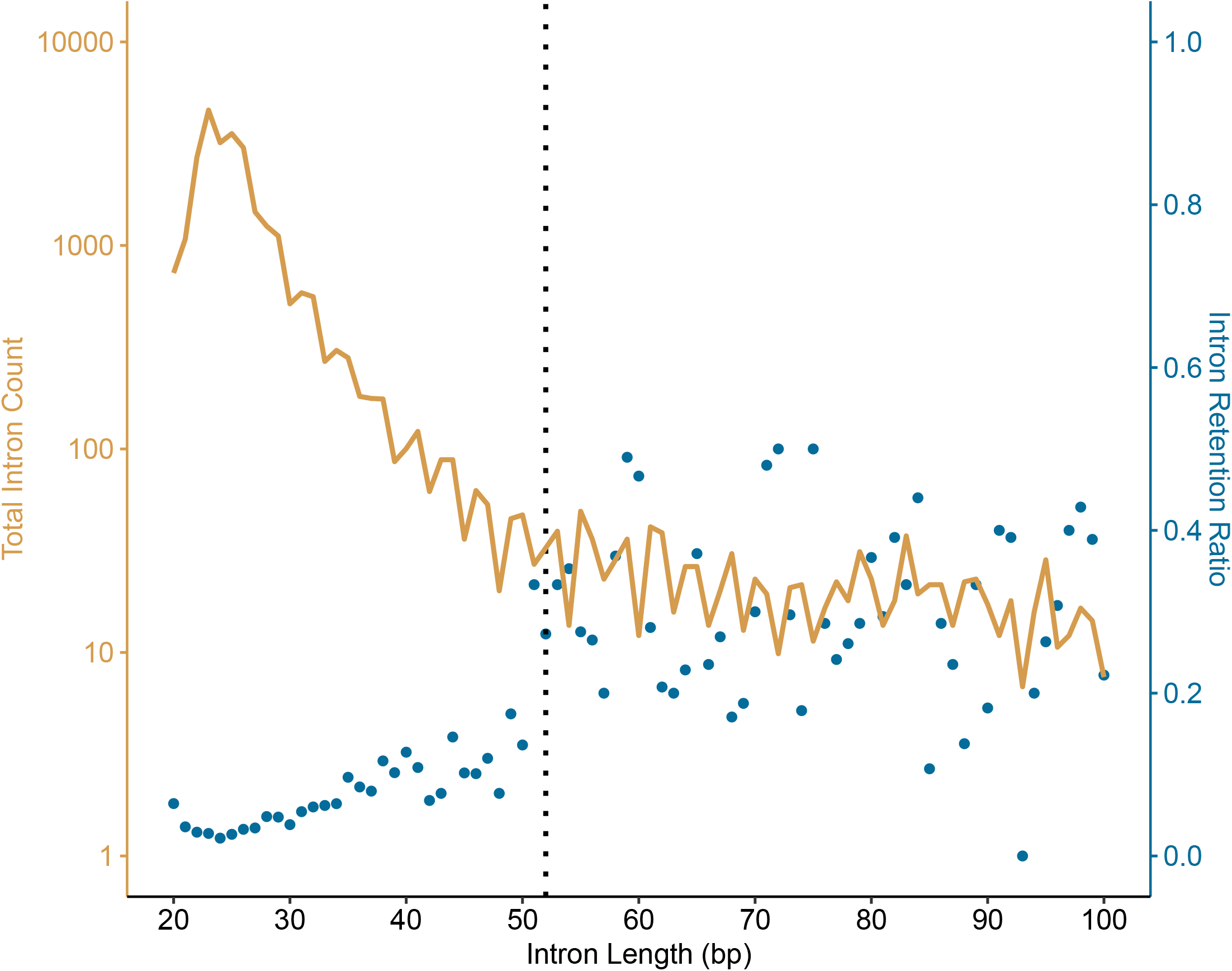
Comparison of intron length profile with detected intron retention (IR). Intron length (bp) is compared against the total number of introns at a range of size from 20 bp to 100 bp (lower end threshold is based on the constraints of bioinformatic software and previous analyses) and shown in light brown (log2 scale). This is contrasted with the intron retention rate of genes with introns that are not excised, shown in blue. A Gompertz function was fitted to the data using non-linear least squares to estimate the intron length greater than which intron retention ratio does not continue to increase (i.e., reaches an asymptote). We computed 95% confidence intervals around the estimated asymptote. The estimated lower confidence bound was then applied as a conservative threshold to identify the intron length (52 bp) at which the intron retention ratio asymptote had been reached (black dotted line). The signal suggests that smaller introns are excised with higher fidelity compared to longer introns which are retained despite there being much fewer longer introns. This suggests intron fidelity is a major source of selection on intron repertoire, consistent with the noisy intron model).

A trend of increased IR rate is present between introns of 25 and 51 bp length (Figure 3). This result is suggestive of a drop in spliceosome efficacy and/or association of weaker splice sites resulting in an increase in IR as introns increase in length [52]. Therefore, our results suggest that IR is a factor among larger *Pb* introns. Conversely, our data also show the population of larger introns has been minimized (number of introns ≥52bp = 1396, 2.66% of the 52,385 total introns which are 100 bp or smaller), suggesting functional selection has played a role in determining the type and distribution of the introns present in the *Pb* 186b genome. These results are consistent with the noisy splicing model [52] being a factor in intron evolutionary dynamics; a model which predicts that if AS is largely a result of splicing errors, the evolutionary dynamics of introns will be driven by selection acting to minimize how introns affect the fidelity of mRNA *trans*lation. For example, AS -particularly IR -will negatively correlate with the length and the expression level of genes [52]. The demonstration that in *Pb* longer introns have higher rates of IR and that longer introns are minimized in number demonstrates an additional outcome of selection on intron repertoire consistent with the noisy splicing model [52]. The strong selection for short introns observed may also be an outcome of the absence of several splicing-related genes which are found in model organisms but which are absent in ciliates [28].

It is interesting to consider why *Paramecia* species can tolerate some intron retention and therefore why *Paramecia* and other ciliates represent a good model system for understanding intron evolutionary dynamics. One factor that may be important is that *Paramecia* have a non-standard genetic code, which means UAG and UAA code for glutamine instead of a stop codons [58–61], while UGA is retained as the only stop codon. As such, the disruptive effect of intron retention on *trans*lation fidelity is limited because fewer codons in the *Paramecia* code for stop codons, therefore reducing the chance that IR would disrupt full-length mRNA *trans*lation. Consistent with this idea, the one ciliate stop codon, UGA, is absent from the common AT rich introns and the possible overlapping codons around the splice site (Figure 2). Also consistent with this result, *Pb* 186b encodes 53,545 introns of 100 bp or less encompassing 1,332,374 possible codons. 39,535 and 111,882 of these possible codons code for UAG and UAA (former stop codons as would be present in the universal genetic code), respectively, while only 13,375 encode the functional UGA stop codon. Again, this suggests that functional stop codons within the *Pb* intron repertoire has been selected against. These dynamics conceivably make *Paramecia* more tolerant to intron expansions and therefore a useful model for understanding evolutionary dynamics, such as IR and subsequent constraints on intron length.

If selection on introns in this manner was indeed a factor, we would also expect introns to locate at a higher frequency towards the 3’ of a gene, thereby reducing the negative impact of IR on coding complete protein domains. To explore this further we looked at the distribution of introns across the entire repertoire of the *Pb* 186b gene repertoire (Figure 4). Briefly, each gene *trans*cript was split into 15 ‘regions’ and introns were mapped to these 15 partitions if their start and stop coordinates were located within the partition. Introns that spanned more than one partition were classified to the partition where the highest percentage of their sequence was located (see Github for R code). This analysis did indeed find a high frequency of introns mapping towards the 3’ of the gene as predicted if selection was acting to minimize the effect of IR. However, this work also identified a high frequency of introns mapping towards the 5’ of the *Pb* genes, suggesting selection is driving introns away from the core regions of the open reading frames in the *Pb* genes. This result is therefore only partially consistent with the idea that selection is shaping the positioning of introns to minimize the effect IR on *trans*lation fidelity, perhaps because IR in *Pb* has indeed a reduced effect on *trans*lation fidelity.

**Figure 4.**
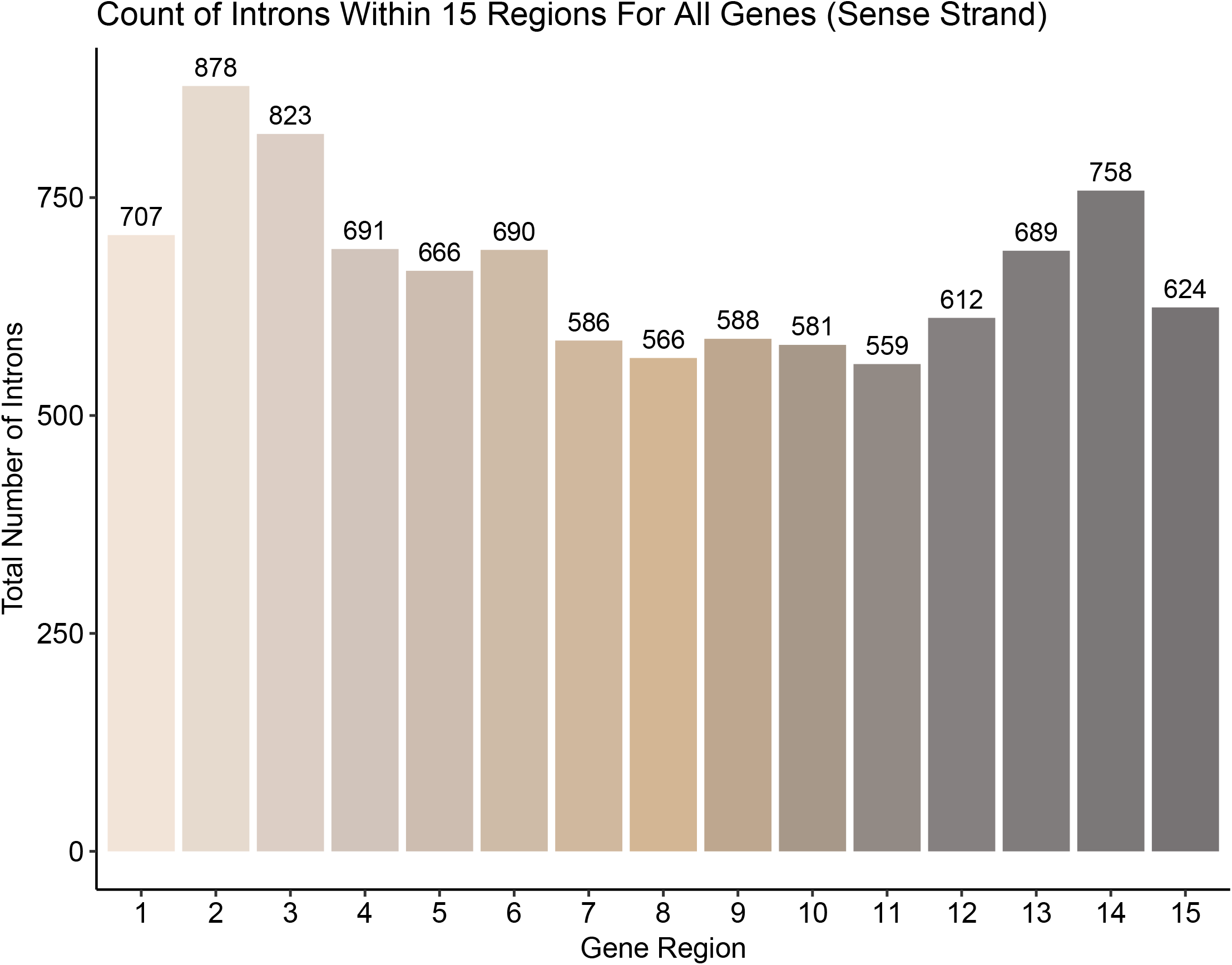
Distribution of where introns locate within the *Pb* repertoire of ORFs. To explore how introns are located within the open reading frames of gene repertoire of *Pb* 186b we divided all genes into 15 equal sections and mapped intron frequency across these 15 sections. These analyses demonstrate that introns are present at higher frequency towards the 5’ and 3’ ends of the ORFs.

## Conclusion

Here we report a highly complete host *Paramecium bursaria* 186b macronuclear genome assembly along with replete *trans*criptome sequencing data using both Illumina and PacBio sequencing methods. This work complements our previous publications which include sRNA sequence datasets [17] and a description of RNAi methodology for targeted knock-down of host-encoded genes [1]. These data are provided to aid genome analysis and to enable the design of forward and reverse genetic experiments for the 186b strain of *P. bursaria*.

Our analyses demonstrate *Pb* 186b possess a large number of tiny introns. To explore intron evolutionary dynamics further we show that intron repertoire in *Pb* 186b is dominated by 23 bp introns and that alternative splicing that leads to intron retention occurs at a higher frequency among longer form introns (>25 bp). This result infers that intron size is under strong functional selection for short introns in the range of 23-25 bp. This pattern of spliceosome selection and higher rates of IR among longer introns is consistent with the noisy splicing model as the primary determinant of intron evolutionary dynamics [52]. We present these data as an aid for further investigation of genome evolution in the *Paramecia*, annotation of *Paramecia* genomes, and the exploration of the rise of endosymbiotic function.

## Data Availability

All sequence reads have been deposited in NCBI GenBank with the BioProject identifier PRJNA659045 for PacBio: SRR12511009, SRR12511010, SRR12511011, Illumina Nova-Seq: SRR12511019, and Illumina ISO-Seq: SRR25546588 (submission in progress). The RNA-Seq *trans*criptome data is available from BioProject PRJNA633103 as condition 1 (SRR11796780, SRR11796779, SRR11796768, SRR11796757), condition 3 (SRR11796746, SRR11796735, SRR11796724, SRR11796713) and condition 5 (SRR11796704, SRR11796703, SRR11796778). The *Pb* 186b genomic assembly will be available at NCBI nuccore (submission in progress) and the predicted protein set will be available from NCBI protein (submission in progress). The genome annotations and other data (including key elements of code and commands) can also be accessed here: https://github.com/guyleonard/paramecium and from Zenodo DOI: 10.5281/zenodo.13240530

## Materials & Methods

### *Pb* 186b Culture Preparation and DNA /RNA extraction

Cultures of *Paramecium bursaria* 186b (CCAP 1660/18) were grown in New Cereal Leaf – Prescott Liquid media (NCL). NCL media was prepared by adding 4.3 mgL^-1^ CaCl_2_.2H_2_O, 1.6 mgL^-1^ KCl, 5.1 mgL^-1^ K_2_HPO_4_, 2.8 mgL^-1^ MgSO_4_.7H_2_O to deionised water. 1 gL^-1^ wheat bran was added, and the solution boiled for 5 minutes. Once cooled, media was filtered once through Whatman Grade 1 filter paper and then through Whatman GF/C glass microfiber filter paper. Filtered NCL media was autoclaved at 121°C for 30 mins to sterilise prior to use.

NCL medium was bacterized with *Klebsiella pneumoniae S*MC and supplemented with 0.8 mgL^-1^ β-sitosterol prior to propagation. *Pb* cells were sub-cultured 1:9 into fresh bacterized NCL media every two months and maintained between 20°C and 23°C with a light-dark (LD) cycle of 12:12h.

*Pb* cells for genomic DNA extractions were concentrated through centrifugation of 10 × 150 ml cultures (10 mins at 800 x *g)* followed by removal of ∼ 80% of the supernatant. Concentrated cultures were then filtered using 15 µm PluriStrainers® and were washed repeatedly with sterile Milli-Q in order to reduce bacterial contamination. Genomic DNA was extracted from pooled cultures using the Qiagen DNeasy Blood and Tissue kit, then the DNA was purified using a Qiagen DNeasy Power Cleanup kit. Long read sequencing was performed using the SMRT Link software version 10.2.0.133434 on a Sequel IIe Pacific Biosciences (PacBio™) device using 6 x SMRT Cells following size selection (> 3 kb) with AMPure PB Beads. The “Run Design Application’’ was set to CLR with default settings.

The resulting PacBio long-read sequence data comprised of six libraries: i) *Pb*1_A05: 5,015,777 reads, 22,472,249,345 bp; ii) *Pb*1_A08: 4,287,200 reads, 18,071,618,742 bp; iii) *Pb*2_A01: 3,985,316 reads, 20,238,646,378 bp; iv) *Pb*2_G10: 3,983,413 reads, 18,192,570,661 bp, v) *Pb*3_A01: 3,019,403 reads, 13,420,246,768 bp, vi) *Pb*3_B01: 3,258,680 reads, 13,820,315,188 bp. In total this includes 23,549,789 reads and 106,215,647,082 bp of sequence

For ISO-Seq, ∼750,000 *Pb* cells in stationary phase were harvested. For NovaSeq, 6 replicate *Pb* cultures were fed for 6 days with HT115 *E. coli ex*pressing non-hit, ‘scramble’ dsRNA [1,17] and ∼500,000 *Pb* cells were harvested. For Iso-Seq and Illumina NovaSeq, *Pb* cells were collected on an 11 μm filter, washed with NCL, and rinsed into 1 mL of TriZol reagent. RNA was extracted using the Zymoprep RNA extraction kit and stored in nuclease-free water at -20 °C. For RNAseq experiments, *P. bursaria* samples were harvested at different points within the Light/Dark cycle (10.5 h into dark cycle “#1”; 6 h into light cycle “#3”; 1.5 h into dark cycle “#5”), as described previously [1].

### Short-Read DNA Sequencing Using an Illumina^™^ NovaSeq

DNA was processed using the Illumina NovaSeq 6000 v1.5 workflow (Illumina^™^) with polyA selection. A 150 bp paired-end library was prepared and resulted in 1,184,917,880 reads total consisting of 177,737,682,000 bp.

### RNA-Seq Sequencing Using Illumina^™^

RNA from a Light/Dark cycle time-course (11 samples: #1A-C,E; #3A,C-E; #5B-D) was prepared for sequencing as described previously [1].

### Iso-Seq Sequencing Using Pacific Biosciences (PacBio^™^)

RNA was processed using the Iso-Seq Express 2.0 workflow (PacBio^™^), targeting *trans*cripts up to 2 kb. Libraries were cleaned using the Express TPK 2.0 (PacBio^™^) and SMRTbell Enzyme Clean-up kit v1 (PacBio^™^), and prepared using the Sequel II Binding Kit 2.2 (PacBio^™^), Sequencing Primer v5 (PacBio^™^) and Sequel II Sequencing Plate v2.0 (PacBio^™^). Sequencing was performed using SMRT link software (10.2.0.133434) on a Sequel IIe machine (PacBio^™^). The “Run Design Application’’ was set to CCS (circular consensus sequencing) with default settings. The resulting PacBio HiFi long-read library consisted of 3,664,630 reads; with total length 8,734,233,994 bp; average length 2,383; and longest length 15,915.

### Long-Read Genome Assembly

The BAM files from the Exeter Sequencing Service [62] were converted to FASTQ using SAMTOOLS v1.15 [63], and then concatenated together. The program LRBinner v2021-06-22 [64] was used to bin the reads using composition and coverage information via a variational auto-enconder. This resulted in twenty-two bins. These bins were then individually assembled using Flye [65] with standard settings. Following these preliminary assemblies, the BUSCO [39,40] tool in ‘auto-lineage’ mode was used to assess each assembly bin for its basic taxonomic profile. This identified three bins (0, 1, & 2) as having strong alveolate signal, three other bins (3, 5 & 7) as having strong bacterial signal (these were not included in any assembly presented here), and all other bins as unclassified (also not included). Binning and subsequent classifications are not exact, and so some of the reads (and subsequent contigs) included may not represent the major taxonomic identity of the bin (i.e., the bins are not 100% clean and therefore may contain some cross ‘contamination’). The raw-reads from the bins with strong alveolate signal (which made up the bulk of the sequencing libraries) were then combined and assembled, again using Flye. The resulting assembly was subsequently polished using Pilon [66] and the Illumina NovaSeq reads (adapter trimmed using FastP [67]), making sure to trim any poly-G tails to account for the 2-colour chemistry of the NovaSeq) for a total of two rounds. Finally, basic cleaning of the resulting contigs (removal of any duplicates) and repeat masking was completed with ‘funannotate clean, sort, & mask’ [50]. This resulted in an assembly with 1,179 contigs, the largest being 395,942 bp, with an N50 of 96,186 bp. Twenty-one bacterial contigs were identified by blasting against NCBI’s ‘nt’ database using BLASTn (as part of the blobtoolkit analysis), and were subsequently removed from the final assembly. A further scaffold was removed on account of being made up almost entirely of repeats. MitoFinder [68] was used to detect presence of mitochondrial contigs in the assembly, using the *Paramecium caudatum* (GenBank: NC_014262) mitochondrial genome assembly as the input. This identified five contigs in total which were removed from the assembly before gene prediction. The five mitochondrial contigs are available from GitHub: https://github.com/guyleonard/paramecium.

### Iso-Seq Assembly

The PacBio software “lima” [69] (to remove barcodes/primers) and the “isoseq3 refine” and “isoseq3 cluster” [51] (to remove polyA tails and cluster *de novo is*oforms respectively) were used to prepare the data from the raw reads (reads: 3,664,630, total length: 8,734,233,994, avg length: 2,383, longest: 15,915). The resulting 97,280 full-length non-concatemer *trans*cripts (total length: 231,414,816 bp) were mapped to the genome assembly using “isoseq3 align” and then isoforms were collapsed with “isoseq collapse”. This produced 44,631 full length *trans*cripts. The final set of *trans*cripts were then *trans*coded to amino acids using the “TransDecoder” [70] pipeline with the Ciliate stop codon usage settings, producing 44,518 peptide sequences. The final set of *trans*cripts were subjected to BUSCO [39,40] analysis and returned Eukaryota ODB10: C:55.7%[S:28.6%,D:27.1%], F:3.5%, M:40.8%, n:255 and Alveolata ODB10: C:87.1%[S:36.8%,D:50.3%], F:0.6%, M:12.3%, n:171. This suggested that we had good coverage of the *trans*criptome. To further assess the coverage of the Iso-Seq *trans*criptome we used pBLAT [71] to search the *trans*cript CDS against the genome, resulting in 44,111 matches at >= 97% identity suggesting 99% coverage (dropping to >= 87% ID returns 100% coverage).

### Genome Annotation

Annotation of the cleaned genome assembly was conducted with GeneMark-ES [72,73] using the Ciliate genetic code table. The gene predictions were then included as input into the FUNANNOTATE [50] pipeline, along with the clustered high-quality Iso-Seq *trans*cripts, and the RNA-Seq *trans*criptome libraries. The funannotate pipeline was directly modified in several places to allow for the alternative genetic code of Ciliates to be used as an option in the underlying programs (e.g., ‘-G Ciliate’ for Trinity, ‘--gcode 6’ for GeneMark-ES v4.71, ‘--stops ATG’, [gcode=6] in tbl2asn amongst other small changes). Functional annotation was provided by multiple databases from InterProScan, (PFAM [74], EGGNOG [75], BUSCO [39,40], Phobius [76], and antiSMASH [[77]) and integrated by funannotate in the ‘other’ mode. Genome annotation fixing (for example to fix erroneously fused genes, and where contigs were split due to adapter contamination) was completed through a process of manual editing of the GFF and FASTA files, the use of the program Liftoff [78] to carry over gene accession names and annotations to the new assembly/gene structures, and followed by the program GAG [79] to help generate the correct file formats for upload to NCBI. Exact commands for the majority of processes in the methods can be found in the GitHub repository (https://github.com/guyleonard/paramecium).

### Intron Identification and Mapping

Introns were extracted from the final set of gene predictions, using the script ‘agat_sp_add_introns.pl’ [80] which adds intron boundaries to the General Feature Format file (GFF) based on the predicted gene structures (exons are present in the GFF but introns are not commonly annotated directly). Their sequences were subsequently extracted using ‘bedtools getfasta’ from the nuclear genome. This was repeated for all other five *Pb* genomes, *P. caudatum* [32] and *P. tetraurelia* [9]. All genomic data and gene prediction GFF files were downloaded from *Paramecium*DB [81]. These were then tallied in R (using phytools [82], stringr [83], dplyr [84], Biostrings [85] and ggplot2 [86]) (Figure 4, see GitHub for code).

### Alternative Splicing Identification

The program IsoQuant [57] was used with a copy of the final gene predictions in GFF format, the nuclear genomic scaffolds and the full length non-concatemer Iso-Seq *trans*cripts mapped to the genome (using HISAT2 [56]) to produce a *trans*cript table of all putative alternate splicing events and their genomic location paired with coverage data and the type of AS event identified.

## Figure legends

**Supplementary Figure 1.** A Gompertz function was applied to estimate the intron length at which the intron retention ratio reaches an asymptote. The red trend line represents values from a Gompertz function which was fitted to the data using non-linear least squares. The blue shaded area represents 95% confidence intervals around the estimated asymptote and the vertical dotted blue line represents the intron length (52 bp) at which the asymptote is reached (applying the lower 95% confidence bound as a threshold).

