## Supplementary figures and images for "*De novo* genome sequence assembly of the RNAi-tractable endosymbiosis model system *Paramecium bursaria* 186b reveals factors shaping intron repertoire"

### Supplementary Figure 1

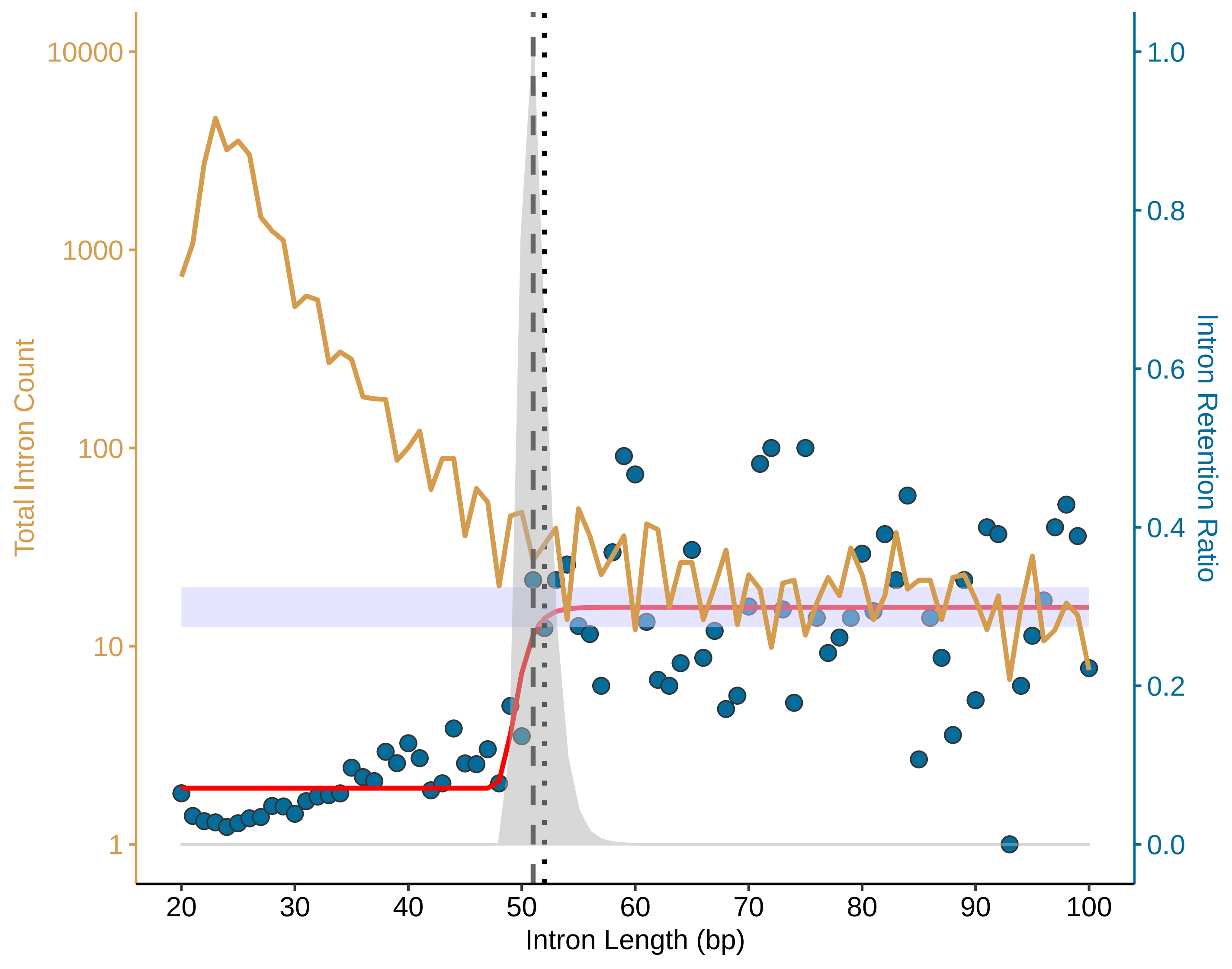
